# Solid optical clearing agents based through-Intact-Skull (TIS) window technique for long-term observation of cortical structure and function in mice

**DOI:** 10.1101/2021.10.02.462855

**Authors:** Dong-Yu Li, Zheng-Wu Hu, Hequn Zhang, Liang Zhu, Yin Liu, Ting-Ting Yu, Jing-Tan Zhu, Wang Xi, Jun Qian, Dan Zhu

## Abstract

In vivo cortical optical imaging needs to overcome the scattering of skull. Compared to the traditional transcranial surgery-based open-skull glass window and thinned-skull preparation, chemical tissue optical clearing techniques can provide a skull-remained optical access to the brain while maintaining its original environment. However, previously demonstrated skull optical clearing windows could only maintain transparency for a couple of hours and hardly capable for high-resolution monitoring of awake animals. Here, we developed a convenient and easy-handling chronic skull optical clearing technique, named “Through-Intact-Skull (TIS) window”, which was compatible with long-term observation at high resolution, and yielded large imaging depth of 900 μm for cortical neurovascular visualization. In addition, our TIS window could monitor neuron activity in awake mice for a long term. Therefore, our bio-compatible and non-invasive TIS window is a new promising approach for intravital brain microscopy with great potential for basic research in neuroscience.

## Introduction

Modern optical imaging techniques provide a powerful tool to observe cortical structure and functions with high resolution and low invasiveness, making it possible to track changes in healthy or diseased brains *in vivo*. ^1-7^ However, the scattering of the turbid skull seriously affects light penetration, limiting imaging quality and depth in the cortex. ^8, 9^ Previous open-skull glass window and thinned-skull window are the two most representative techniques that had been widely used in neuroscience.^10, 11^ However, the thinned-skull window requires high skills to grinding the skull to ∼20 μm, which is also inconvenient for long-term repeatable observation. The open-skull window allows for long-term imaging, but not suitable for immediate imaging for brain recovery. In addition, these surgery operations could change intracranial pressure, trigger a series of inflammatory reactions, thus altering the native state of the brain. ^12^

Compared to the open-skull glass window and thinned skull window, the recently developed optical clearing technique provides an easy-handling, noninvasive and efficient skull window for brain optical imaging.^13, 14^ By applying biocompatible reagents to the surface of skull, the scattered particles in skull, such as collagen, lipids and calcium, could be removed, and the refractive index (RI) of the skull becomes uniform, making the skull transparent. In this way, the brain could be visualized with an intact skull, avoiding potential undesirable damage.^15-17^ With the assistance of the skull optical clearing window, deep-tissue cortical vascular structure (around 300-μm depth) observation and short-term brain neurodevelopment monitoring (with Dendritic spines resolved) have been realized. ^13, 14^

Despite the above key advantages of the skull optical clearing window, it still has limitations in long-term repeatable observation. The skull could keep transparent only when it is well covered by optical clearing agents. Since the reported *in vivo* skull optical clearing agents are liquids,^13, 14, 18^ the optical clearing treatment to the skull is needed every time before imaging, making it rather inconvenient. In addition, when using water-immersion objective for high-resolution imaging, a plastic wrap was usually used to isolated water and liquid optical clearing agents.^13, 14^ In this case, such interfaces (agents/ plastic wrap and plastic wrap/water) are easily interrupted by animal’s slight moving, leading to a decreased imaging quality. Consequently, the previous optical clearing skull windows are hard to observe cortical fine structure or activity in awake animal. Therefore, we set out to develop a new solid-state skull optical clearing method that enables long-term intact skull transparent for a long time, thus permitting longitudinal brain visualization over 3 weeks.

In this work, we established a chronic skull optical clearing window in mice through collagen dissociation, degreasing, RI matching, and curing, after which the skull could keep transparent in weeks, while the existed *in vivo* skull optical clearing methods were only used to make the skull transparent for one-time observation, which usually last for hours. ^13-17^ We named this novel technique “Through-Intact-Skull window (TIS window)”. We demonstrate the efficacy of our technique for cortical structure and function imaging for a range of optical imaging modalities, including laser speckle contrast imaging (LSCI), hyper-spectral imaging (HSI), two-photon microscopy, and three-photon microscopy. The results showed that the TIS window significantly improved the image quality for each imaging method. Specifically, through the TIS window, both apical dendritic spines located in the superficial cortex (∼ 150 μm) and the neuronal cell bodies located in the deep cortex (>800 μm) could be clearly observed by multi-photon microscopy for weeks. In addition, the TIS window also allowed us to monitor stimuli evoked neural responses for weeks in awake mice. A series of *in vivo* and *in vitro* experiments indicated that both microglia and astrocyte, and neutrophil did not show enhanced activities under the TIS window, corroborating its biocompatibility. Therefore, the newly established skull-remained and inflammatory-free TIS window remarkably increased the transparency of mice skull for weeks, during which period deep-tissue and synaptic-resolution imaging could be performed anytime, even in their awake state. Our novel technology holds great potential in the investigation of neuroscience combine with many optical methods.

## Results

### TIS window establishment

We designed a long-term TIS window to make the skull transparent without any transcranial surgery, as shown in Fig. 1a and 1b. Firstly, we treated skull with S1 to make collagen in the skull dissolute.^19, 20^ Secondly, we used sodium S2 to make skull ulteriorly uniform.^21^ Thirdly, we used S3 to perform refractive index matching. Fourthly, we added a coverslip to S3 to guarantee its top surface flatness which is important to minimize optical aberrations. Finally, we introduced light irradiation to solidify S3, bounding the skull and the coverslip. In this case, the TIS window is compatible for both air-immersed lenses or water-immersed objectives.

**Figure 1.**
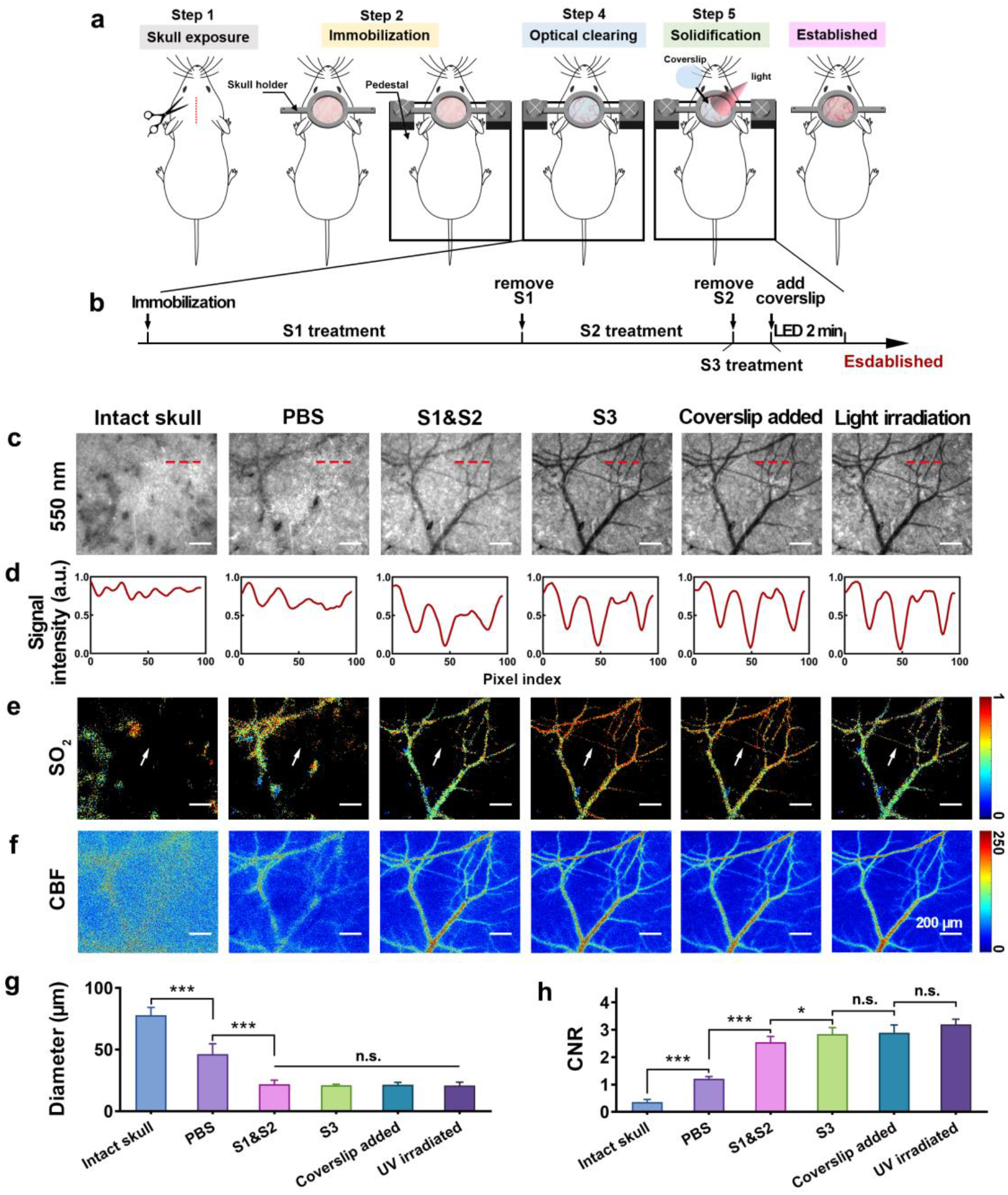
The Through-Intact-Skull (TIS) technique. **(a)** Illustration of the TIS window establishment. **(b)** Details of step 4 and step 5 in (a). **(c)** Typical 550-nm reflectance maps (at 550 nm) of cortical vessels under different conditions. **(d)** Signal intensity along the dashed lines in (c). **(e)** SO_2_ images of the same ROI in (c). The white arrows indicate a blood vessel that could not be observed by SO_2_ imaging before S3 treatment. **(f)** Cerebral blood flow (CBF) images of the same ROI in (a). **(g)** Bar graph of minimum resolvable vascular diameter for mice under different conditions based on CBF images. **(h)** Contrast-to-noise ratio (CNR) under different conditions based on LSCI images. *p<0.05, ***p<0.001, and N.S.: no significant difference.

Before evaluating the TIS window *in vivo*, we performed *ex vivo* experiments to examine the effect of the reagents involved in the establishment of the TIS window. After S1 and S2 treatment, the transmittance of the skull in the band of 500-1000 nm was increased by 10 times from originally less than 5%, to about 50% after S3 treatment and curing, making the grid paper below the skull visible. In addition, the 1951 United States Air Force (USAF) resolution test target showed it was not visible through the intact skull, while the resolution increased with S1, S2, S3 and curing treatments (250 μm vs. 110 μm vs. 12.5 μm vs. 7.81 μ).

To test the efficacy of the TIS window *in vivo*, bright field imaging at 550 nm, HSI for oxygen saturation (SO_2_) imaging, as well as LSCI for cerebral blood flow (CBF) imaging were firstly performed during the core steps of window establishment. As shown in Fig. 1c and 1d, before optical clearing treatment, no vasculature could be observed under 550 nm, which didn’t change much with PBS treatment. However, during the optical clearing processing, even small cortical blood vessels became visible with promising contrast. The results of HSI and LSCI both showed that the functional information of cortical blood vessels became clearer during the skull optical clearing procedure (Fig. 1e and 1f). In addition, the LSCI images were quantitatively analyzed. As shown in Fig. 1g, through the intact skull, only blood vessels with diameters as large as 77.8 μm could be distinguished, and this number decreased down to 46.3 μm when the skull was covered with PBS. However, the minimum resolvable vascular diameter remarkably reduced during TIS window establishment (20.8 μm). The above numbers were determined by the smallest observable blood vessel in each blood flow images. Moreover, the contrast-to-noise ratio (CNR) got increasingly better with the treatment of S1&S2 and S3 (Fig. 1h). Here, the resolve power was still limited by the imaging device, which just used a 2× objective and an industrial CCD (512 pixels×512 pixels; pixel size: 6.45 μm×6.45 μm).

### TIS window for multi-photon microscopy imaging

Next, the imaging resolution and imaging depth through the TIS window were evaluated via multi-photon microscopy for recording the cortical vasculature and neurons. As shown in Fig. 2(a1), dual-channel neurovascular two-photon microscopy was performed through a dual-hemispheric TIS window. Both large blood vessels and tiny microvasculature in the cortex could be clearly visualized. For the open-skull window, it is inconvenient to observe cortical areas around the sutura, because removing the skull above these areas can cause massive bleeding. On the contrary, with the TIS window, it was easy to view the areas around the sutura without worrying about bleeding injury (Fig. 2(a2-a4)), and the dendritic spines can be clearly observed (Fig. 2(a5-a10)).

**Figure 2.**
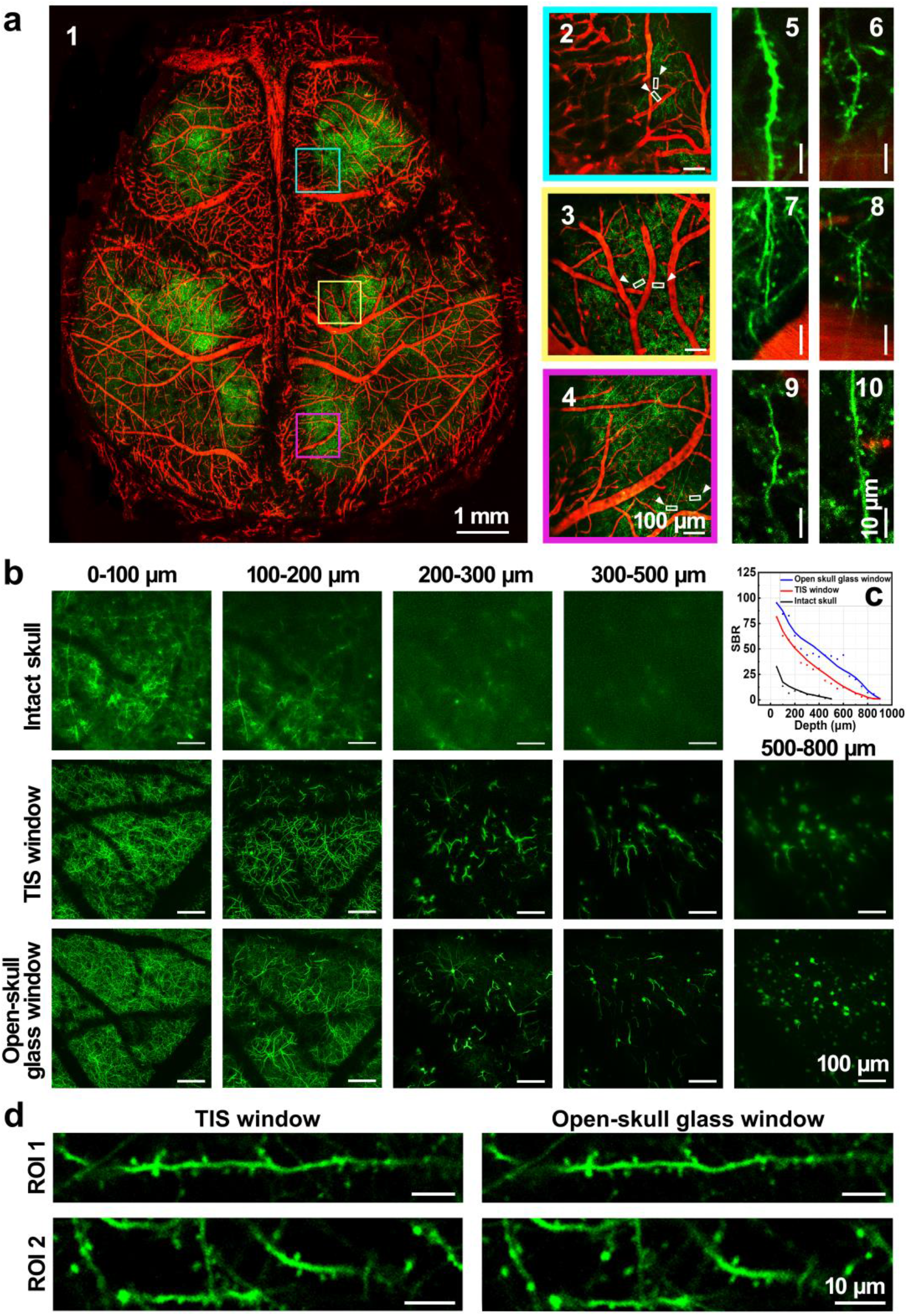
TIS window for two-photon microscopy. **(a1)** A large-field and dual-channel two-photon microscopic image for bilateral hemispheric cortical blood vessels (in red, Texas red) and neurons (in green, EGFP) after the establishment of the TIS window (collected with a 4× objective). **(a2-a4)** High magnification images of the corresponding color frames in (a1) near the sutura (collected with a 20× objective). (a5-a10) High magnification (zoom in) images of the white frames (white arrow indicated) in (a2)-(a4) correspondingly. (20× objective, zoom 4). **(b)** Typical images of two-photon microscopy for neurons at various depths before, after the establishment of the TIS window, and after the skull was removed. **(c)** The SBRs of the images at various imaging depth under different conditions. The SBR was determined as follows: for each image of certain imaging depth, representative dendrites were chosen and the observed intensity of them was recorded as the signal, and the average intensity around the dendrites was recorded as the background, therefore the SBR could be calculated. **(d)** The comparison of TIS window and open-skull glass window on dendritic spine imaging. The excitation wavelength was 920 nm.

We then used two-photon microscopy to visualize neuronal morphology and structure and to compare the image quality and achievable imaging depth of the initial state skull, TIS window establishment and skull removal. As shown in Fig. 2b, through the original skull, details of the apical dendrites were hard to resolve, although located on the very surface of the cortex. Axons and cell bodies in deep tissue were obscured by high background signals and the profiles of the neurons were hardly distinguishable at 400-μm depth. However, after the TIS window was established, the apical dendrites, axons and cell bodies in the cortex could be observed clearly with two-photon microscopy, and the final imaging depth could easily reach over 500 μm. Although the imaging contrast through TIS window was slightly lower in large depth than that through open-skull glass window, it was much more promising than that through intact skull. (Fig. 2c) In addition, compared with the skull opened imaging, the resolution at shallow cortex through TIS window was similar in superficial layer (Fig. 2d), with a slightly lower resolution in deep layer over 500 μm.

Furthermore, two previous skull optical clearing window techniques were also performed for comparison. As shown in Fig. 3 and 4, compared to SOCW^13^ and USOCA^14^, both the resolve power and imaging contrast through TIS window were remarkably better.

**Figure 3.**
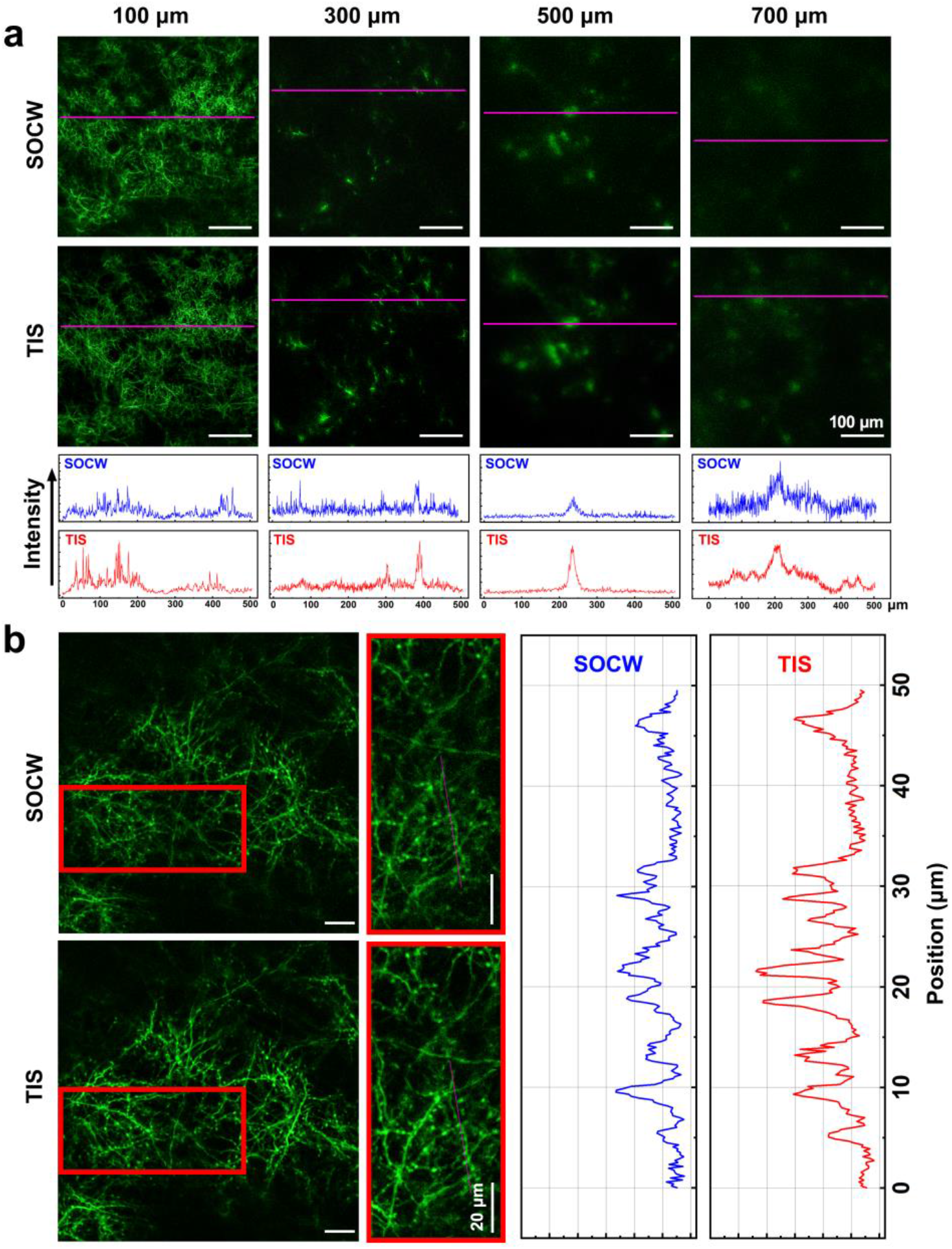
Comparison between SOCW and TIS technique via two-photon microscopy imaging of cortical neurons. (a) Two-photon images at various depths. The plots represent the intensity distributions on the lines. (b) zoom-in images at 150-μm depth. The plots represent the intensity distributions on the lines. The plots showed that the full width at half maximum of the imaged structure was smaller, and the signal intensity was stronger through TIS window, indicating a better imaging quality.

**Figure 4.**
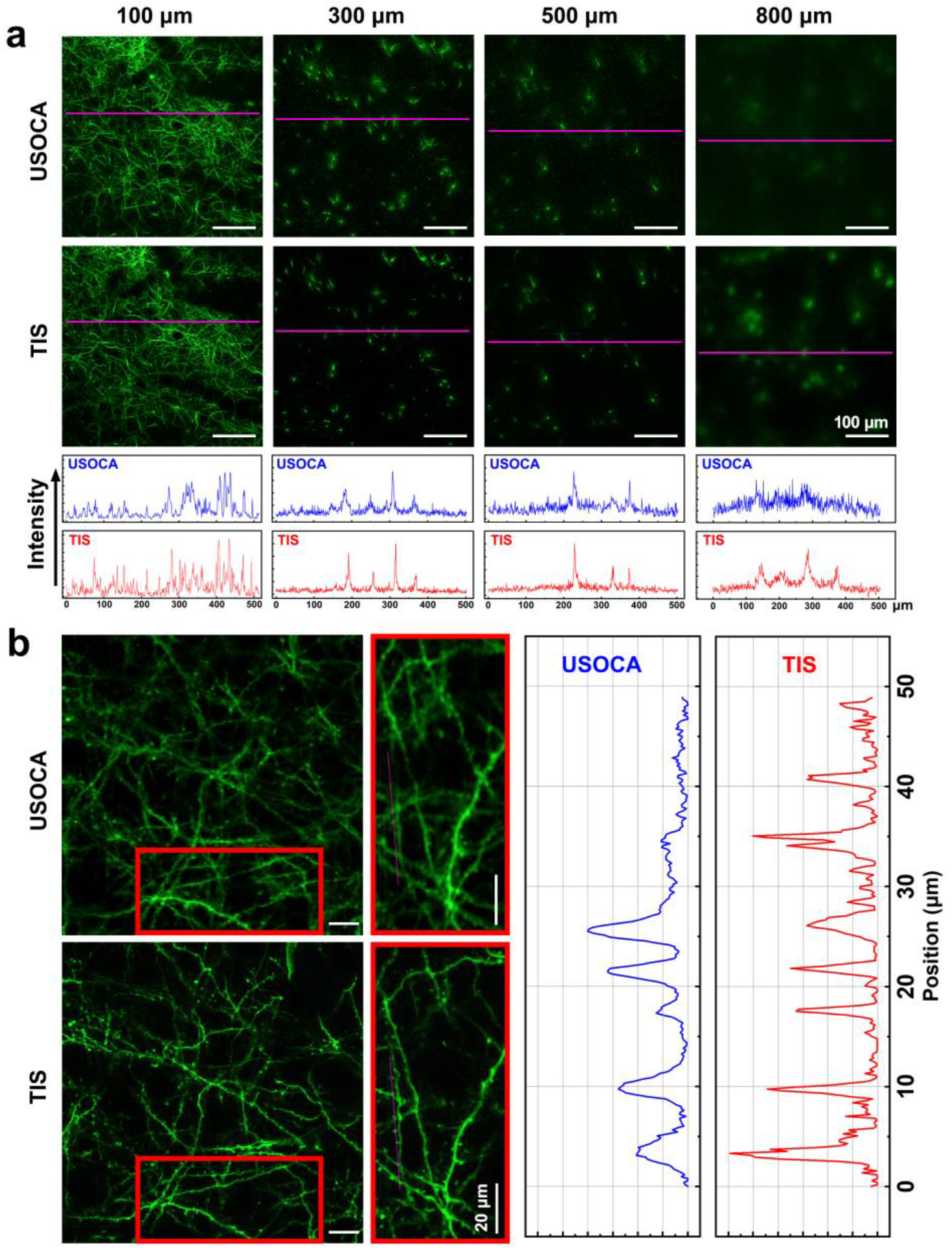
Comparison between USOCA and TIS technique via two-photon microscopy imaging of cortical neurons. (a) Two-photon images at various depths. The plots represent the intensity distributions on the lines. (b) zoom-in images at 150-μm depth. The plots represent the intensity distributions on the lines. The plots showed that the full width at half maximum of the imaged structure was smaller, and the signal intensity was stronger through TIS window, indicating a better imaging quality.

To push the achievable imaging depth, we also evaluated our TIS window with three-photon microscopy which holds a key advantage of less out-of-focus background and therefore a better signal-to-background ratio (SBR) at large tissue depths compared to two-photon microscopy. Although three-photon microscopy uses longer excitation wavelength than two-photon microscopy, thus suffers less scattering of tissue, the signal light in short wavelength was still affected by tissue scattering. Therefore, without additional treatment, the through-skull three-photon neural imaging depth was around 400 μm,^22^ much less than that when the skull was removed.^23^ Therefore, it is valuable to evaluate if TIS window could be combined with three-photon microscopy to reach deeper tissue through intact skull. Since two-photon imaging and three-photon imaging utilize different excitation wavelengths, it is important to test the TIS compatibility with the longer wavelengths needed for three-photon imaging.^24^ As shown in Fig. 5a, through the turbid skull, three-photon microscopy permitted us to observe some information at a certain depth, but few signals below 400 μm. On the contrary, through the TIS window, more synapses in the superficial layer could be observed and abundant cell bodies came out with promising imaging quality in the deep layer (400-800 μm), which could not be monitored without TIS window (Fig. 5b and 5c). The dendrites could still be distinguished even at depth of 800-900 μm. In addition, the 3D shape of cell bodies could be finely reconstructed (Fig. 5d). in addition, the SBR at different imaging depths was collected (Fig. 5e). The result showed that, through the turbid skull, the SBR at 100 μm was 20, but decayed rapidly to ∼1 at 500 μm. Through the TIS window, the SBR at each depth was significantly higher than that through the turbid skull. It was ∼60 at 100 μm, and still near 20 at 700 μm.

**Figure 5.**
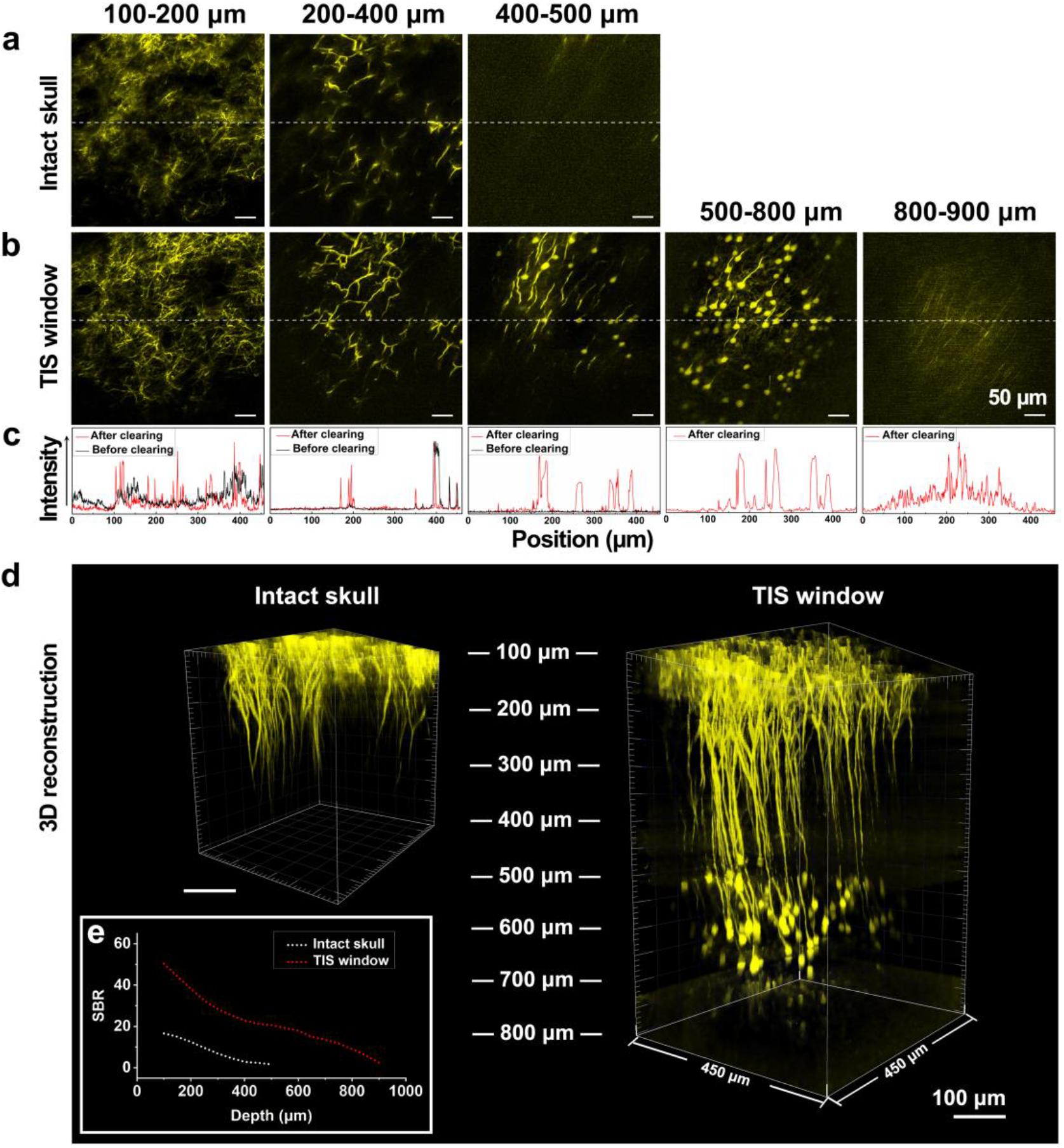
TIS window for three-photon microscopy. **(a-b)** Representative images of three-photon microscopy for neurons in a Thy1-YFP mouse at 1300nm excitation wavelength and at various depths before (a) and after (b) establishment of TIS window. **(c)** Signal intensity along the dashed lines in (a) and (b). **(d)** 3D reconstruction image before and after the establishment of TIS window. It was only achievable above 400 μm through intact skull. **(e)** SBR at each depth before and after TIS window establishment.

### TIS window for long-term cortical vascular observation

To evaluate the ability of the TIS window to maintain skull transparency, we next performed long-term monitoring of vasculature in the superficial and deep cortex via two-photon microscopy imaging and LSCI. As shown in Fig. 6a, through the intact, turbid skull, only large vasculature above 100-μm depth could be clearly resolved, while relatively small vessels remained blurry. On the contrary, with the assistance of the TIS window, abundant tiny microvasculature could be monitored with high contrast at a depth as large as 700 μm. Even 10 days after the TIS window establishment, with two-photon microscopy the tiny microvasculature could be easily discerned, and the imaging depth remained as large as 900 μm.

**Figure 6.**
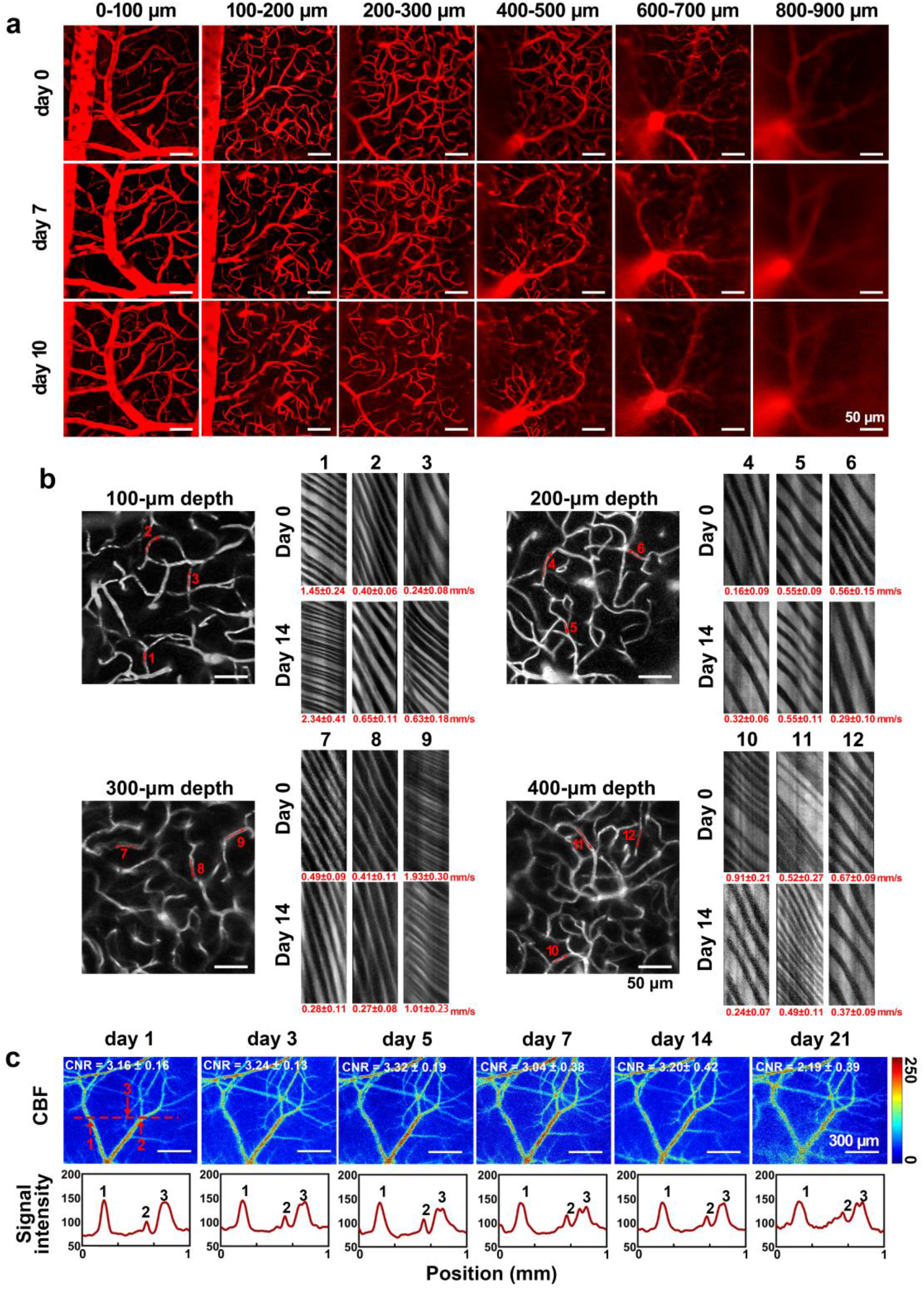
TIS window for long-term cortical blood flow monitoring. **(a)** Representative cortical vascular images of two-photon imaging at various depths on day 0 (the day TIS window was established), day 7 and day 10 after TIS window establishment. (b) Representative two-photon line scanning images for blood vessels speed measurement at various depths on day 0 and day 14 after TIS window establishment (red number in depth image indicated line scan position, corresponding to the line scanning images). **(c)** Representative cortical vascular LSCI images on day 1 (the next day of TIS window establishment), day 3, day 5, day 7, day 14 and day 21 after TIS window establishment. The plots represent the signal intensity along the dashed lines in the LSCI image on each day.

In addition to structural observation, we also demonstrate the capability of our TIS to aid in monitoring vascular function, such as blood flow velocity. As shown in Fig. 6b, under the TIS window, various blood flow profiles could be measured at different depths. Figure 6c showed the results of long-term LSCI observation during 1-21 days after the establishment of the TIS window. Over the entire 2-week period, the skull remained transparent, and the CNR of the cortical blood flow imaging quality maintained at a high level. In the 3rd week, the CNR of LSCI was slightly reduced, but all blood vessels who were monitored during the first two weeks could still be distinguished. This indicated that the vascular function could be tracked with various imaging methods under the TIS window over 14 days.

### TIS window for long-term neural imaging

We next applied the TIS window technique for long-term monitoring of neurons at synaptic resolution in deep tissue using multi-photon microscopy. As shown in Fig. 7a, the imaging quality and imaging depth of two-photon deep-tissue microscopy had negligible decrease over time, and axons can be tracked down to a depth of 800 μm during 2 weeks after TIS window establishment.

**Figure 7.**
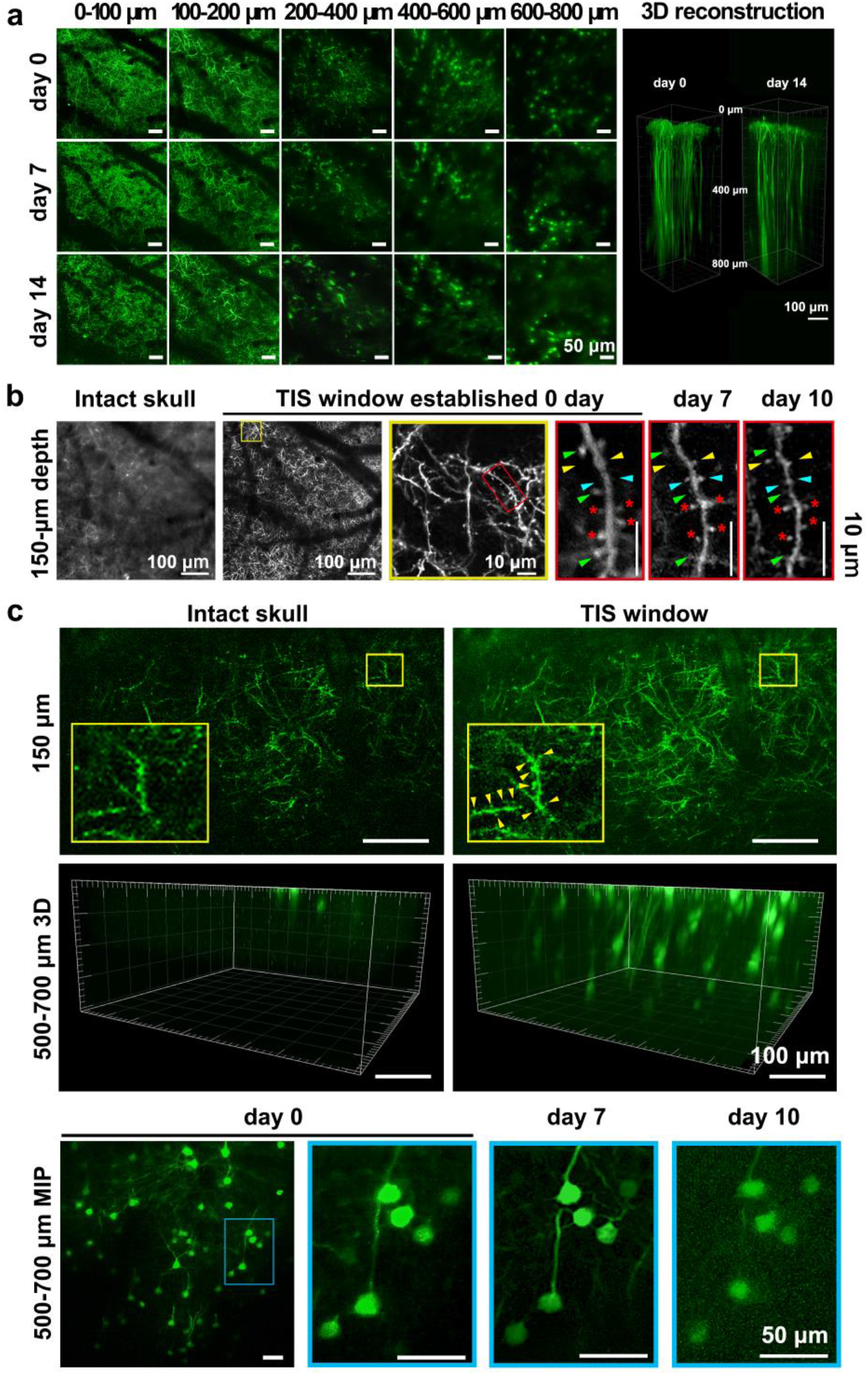
TIS window for long-term neural monitoring. **(a)** Representative images of two-photon cortical neural imaging in a Thy1-EGFP mouse at various depths on day 0, day 7 and day 14 after TIS window establishment. **(b)** long-term two-photon observation of dendritic spines through TIS window at 150-μm depth. Green arrows show spines that were eliminated during 10 days; yellow arrows show spines that were formed during 10 days; cyan arrows show spines that were formed and then eliminated during 10 days; asterisks show spines that keep existing during 10 days. **(c)** Representative images of three-photon microscopy for synaptic imaging and neural cell body long-time monitoring in deep tissue. Yellow arrows point to the spines those could be distinguished, according to the TIS window technique. The excitation wavelength was 920 nm and 1300 nm for two-photon microscopy and three-photon microscopy, respectively. MIP: maximum projection.

The morphology and number of dendritic spines undergo natural changes over time, and the abnormal dynamics of dendritic spines could reflect nerve diseases. Therefore, it is important to investigate the longitudinal dynamics of dendritic spines in vivo. Thus, the capability of the TIS window to perform long-term observation of dendritic spines was evaluated. As shown in Fig. 7b, through the intact turbid skull, two-photon microscopy could not provide sufficient resolution to observe dendrites at 150-μm depth, nor dendritic spines. On the contrary, with the TIS window technique, the long-term dynamics of spines could be observed reliably. The formation and elimination of dendritic spines have been clearly observed for 10 days. This is a promising result that shows our techniques potential to monitor synaptic dynamics and their role in motor memories and neuronal circuit establishment, as well as brain dysfunctions. For instance, within such a period, the synaptic plasticity was observed to be very different in the Parkinson’s disease (PD) ^25, 26^.

Figure 7c showed representative images of three-photon microscopy in which dendritic spines could be visualized, but with lower resolution than two-photon microscopy (Fig. 7b), probably due to the slightly increased PSF at 1300 nm compared to 920-nm based two-photon microscopy. Still, three-photon microscopy holds the advantage of high resolution and high SBR, and thus can be utilized to track neural cell bodies in deep tissue. As shown in Fig. 7c, the cell bodies at 500-700-μm depth were clearly observed through the TIS window for 10 days, while it could be observed that with two-photon microscopy, the cell bodies could not be distinguished in the 3D reconstruction (Fig. 7a). Therefore, through the TIS window, long-term fine visualization of neurons from surface to deep cortex could be performed with the combination of two-photon and three-photon microscopy.

### TIS window for long-term neural functional imaging in awake mice

We then applied the TIS window technique to monitoring the calcium mediated neural activity in an awake mouse. As shown in Fig. 8a, the TIS window was established right after Gcamp6s virus injection for 14 days. Figure 8b show an ROI that contains 14 neuron dendrites during the mouse paw moving. Through the TIS window, the spontaneous activity of each dendrite could be clearly monitored while the mouse was awake. As shown in the results, at the quick scanning imaging mode (24 fps), the TIS window allow us to distinguish dendrites around 2 μm.

**Figure 8.**
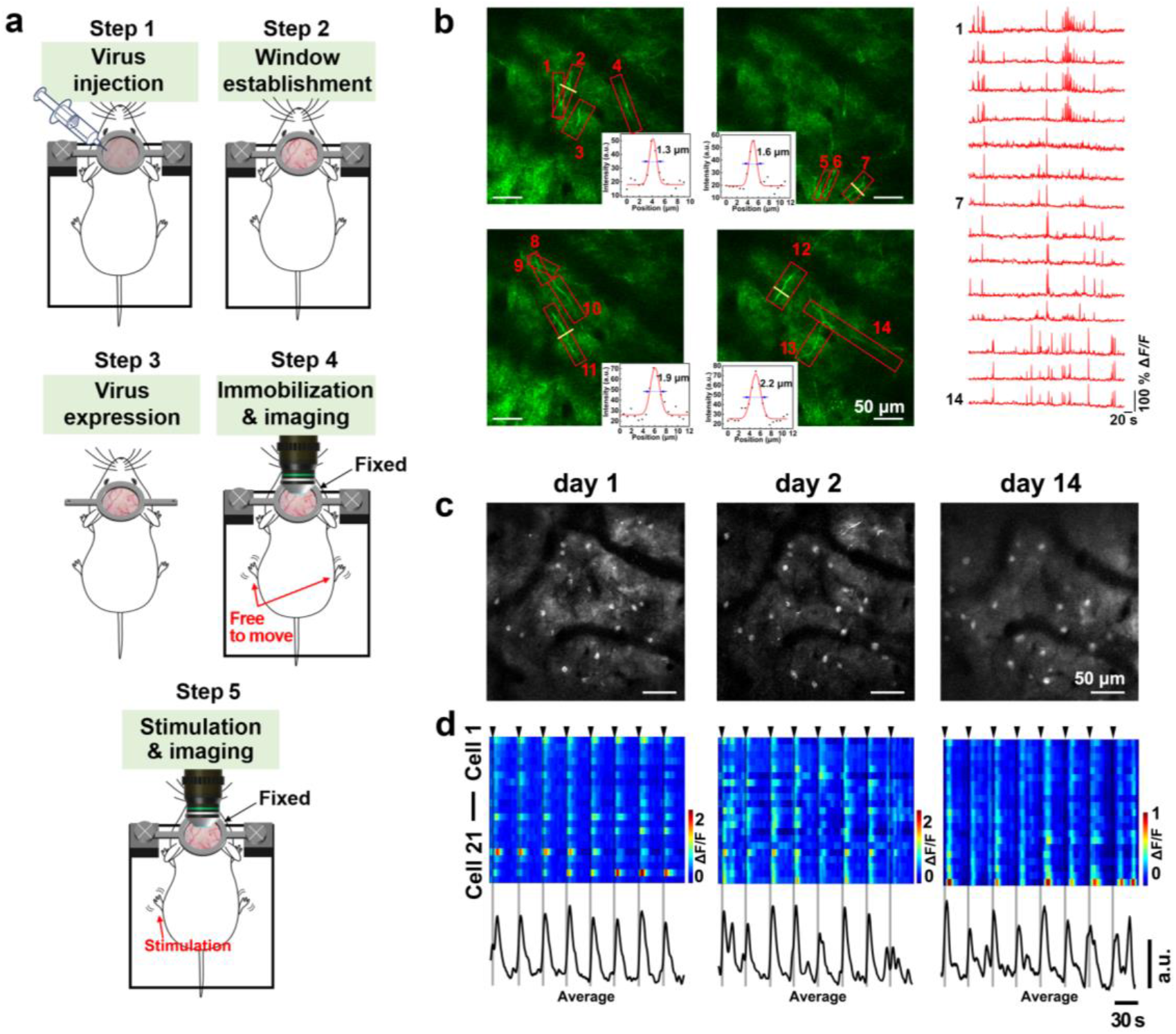
TIS window for neural activity monitoring in awake mice. **(a)** Illustration of TIS windows for GCaMP6s imaging in awake mice. **(b)** Representative two-photon image of dendrites marked with GCaMP6s, and their spontaneous activities. The red numbers represent the individual dendrites and the corresponding dynamics of the GCaMP6s activity. The inset plots represent the intensity distribution of the yellow lines. **(c)** Representative two-photon images of neural bodies marked with GCaMP6s on day 1 (the 11^th^ day after TIS window establishment), day 2 and day 14. **(d)** Neuronal responses of 21 cells (up) and the average response (down) on day 1, and 2 and day 14 (black arrow and gray bar indicate ES stimuli).

Electro-stimulation (ES) for the forepaw was used to evoke neuronal activity in the somatosensory cortex. Figure 8c show an ROI that contains 21 neurons. Through the TIS window, most of the neurons showed observable responses to the ES for day 1 to day 14 under the TIS window. (Fig. 8c and 8d) This data suggests that the same neuron population activity can be tracked for weeks during the TIS window under two-photon imaging.

### TIS window for optogenetics in awake mice

In addition to optical observation, we further evaluated the capability of TIS window for optogenetic manipulation in wake mice. As shown in Fig. 9, we expressed Thy1-ChrimsonR-tdTomato (red channel) and Thy1-GCaMP6s (green channel) in neurons in somatosensory cortex. When the 590nm laser stimuli on the TIS window, the neurons showed active responses after the stimuli. These results indicated that TIS window also holds great potential in optogenetic manipulation of the neural circuits with imaging methods.

**Figure 9.**
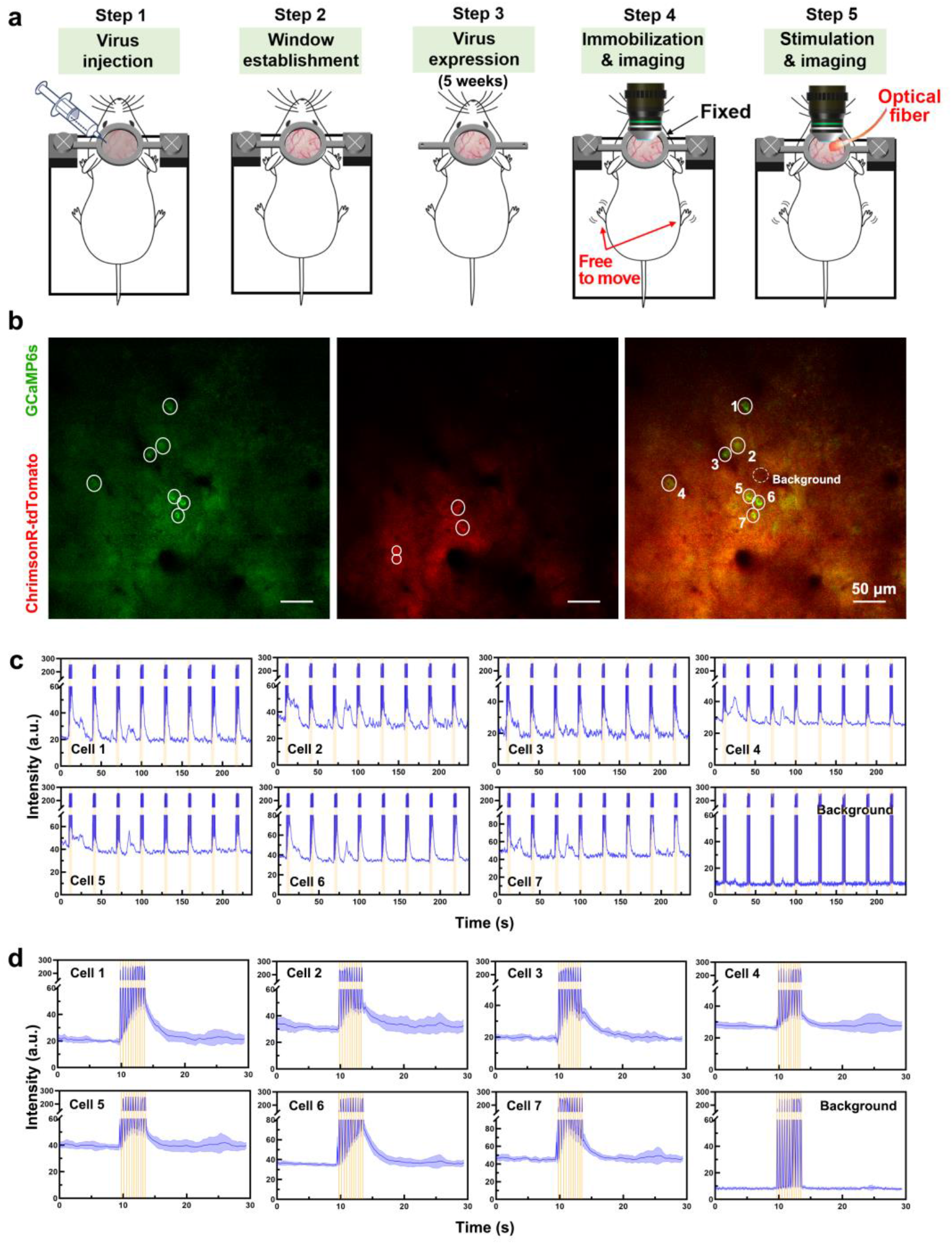
**(a)** Illustration of TIS windows for optogenetic manipulation and imaging in awake mice. **(b)** The representative image of neurons marked by GCaMP6s and ChrisonR-tdTomato. **(c)** Quantitative analysis of neural response to 590-nm laser stimulation (10ms /2.5Hz 4s) for 8 trails. **(d)** Average neural response of the 8 trails in (c).

### Biosafety assessment of TIS window

While the long-term repeated imaging of neurons and blood vessels in the same positions without morphological changes as shown in above Figures, suggesting the absence of severe damage or toxicity due to our TIS window, we additionally performed both *in vivo* and *ex vivo* experiments to further assess the biosafety of our method. First, we monitored microglia *in vivo* at 1 hour and 48 hours after TIS window establishment. The microglia remained at the same position with an inactive state (Fig. 10a), which indicates that the TIS window doesn’t lead to microglia activation.

**Figure 10.**
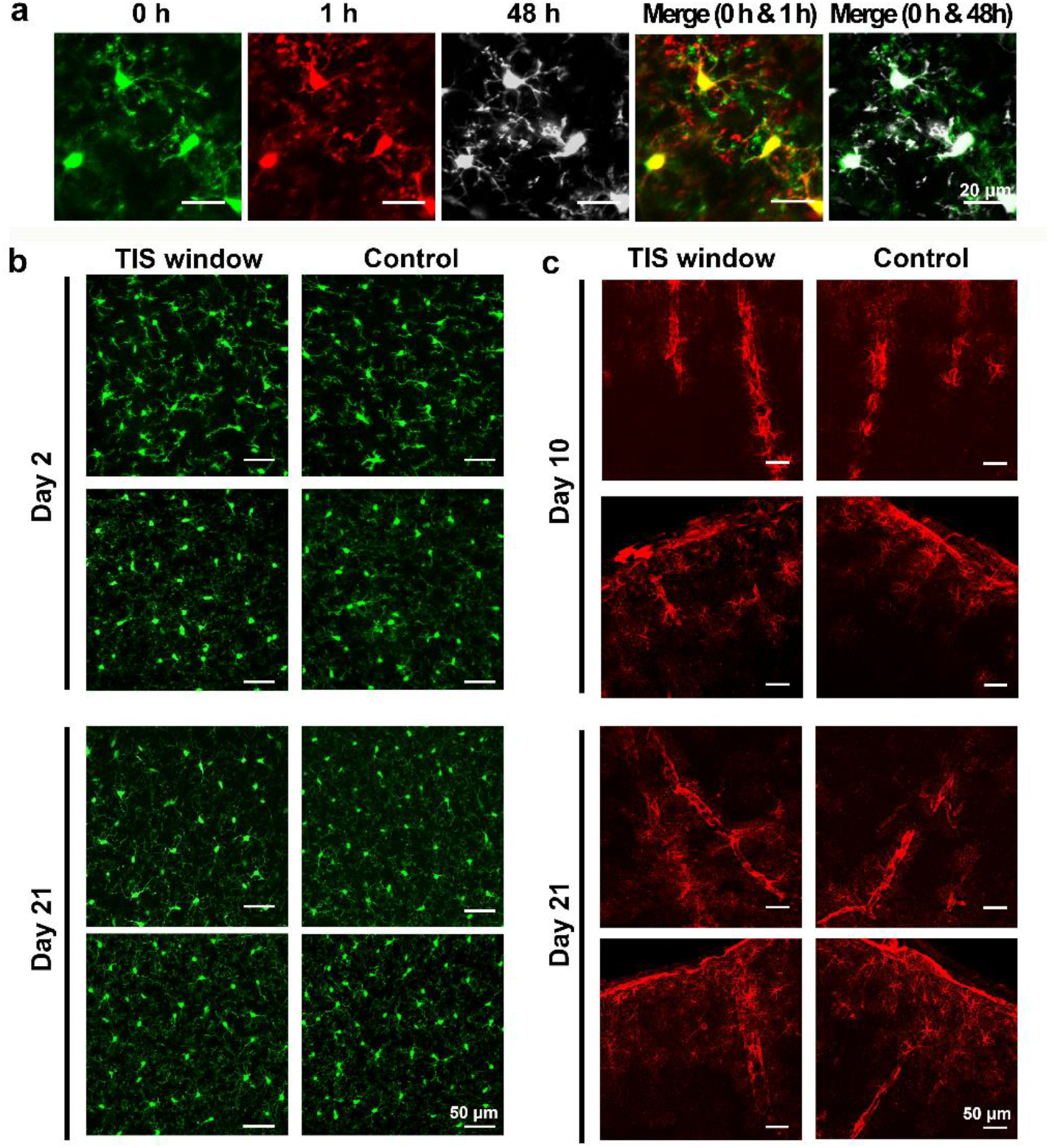
(a) Representative *in vivo* microglia images immediately, 1 hour and 2 days after TIS window establishment. (b) Representative microglia images in brain slices 2 days and 21 days after TIS window establishment. (c) Representative GFAP expression in astrocytes in brain slices 10 days and 21 days after TIS window establishment.

Next, we perform a series of *ex vivo* experiments. As shown in Fig. 10b, the microglia in both the TIS window and the control hemisphere remained in a non-active state with a highly branched morphology, and the volume and the ellipticity had no significant difference in the two parts of the hemisphere. In addition, the GFAP immunofluorescence images showed similar intensity and distribution for both sides (Fig. 10c), which means that the astrocytes were not activated. Furthermore, the H&E staining imaging suggested that, compared with contralateral, there was no difference in the brain parenchyma under the TIS window (Fig. 11), indicating the TIS window won’t cause inflammation.

**Figure 11.**
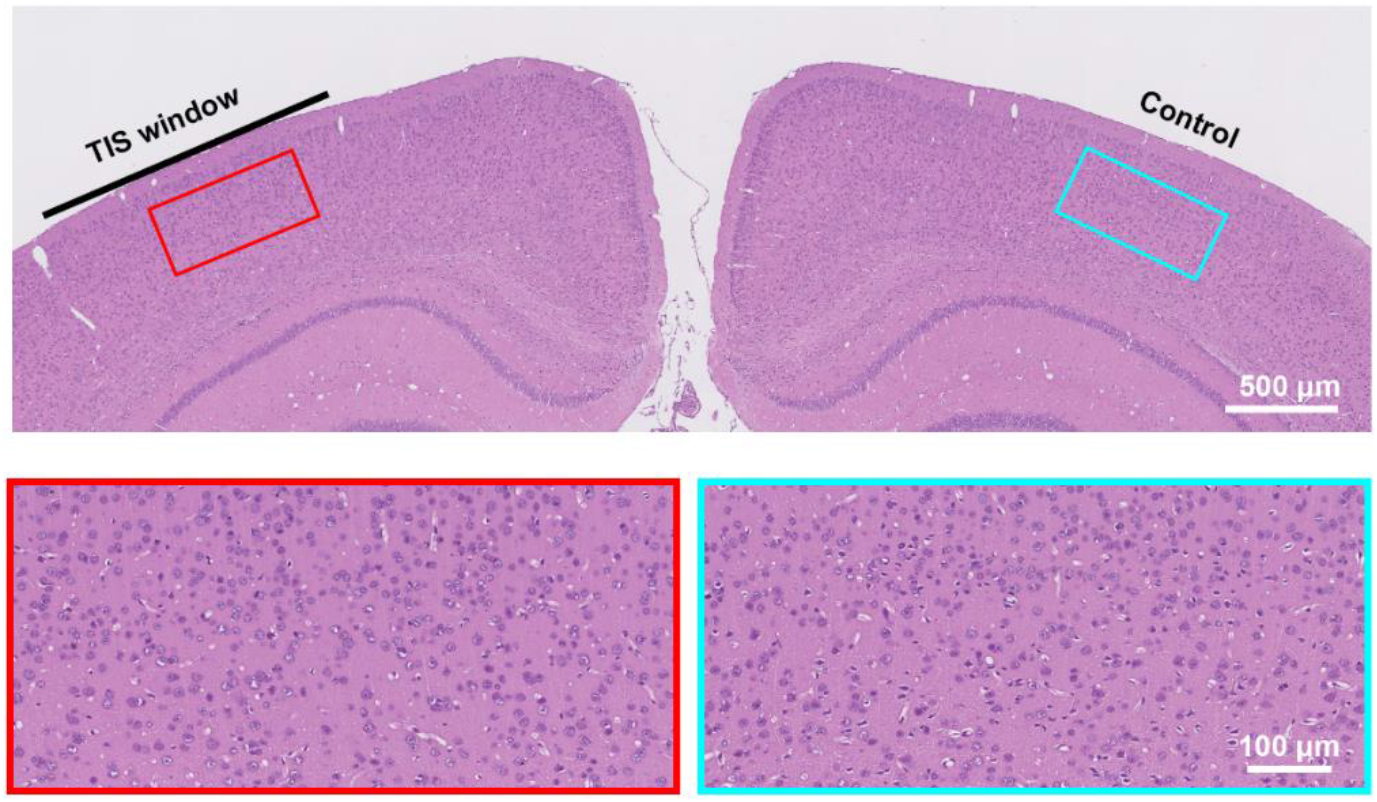
Representative images of H&E stained brain slice 2 days after TIS window. establishment.

## Discussion

In this work, we established a through-intact-skull window with several optical imaging methods *in vivo*, making the skull stay transparent over weeks for long-term cortical structure and function imaging. This TIS window allowed us to obtain long-term cortical imaging without significant resolution loss or imaging depth decrease. For instance, the dynamic of dendritic spines of neurons and microvasculature could be observed for weeks in an awake mouse. In addition, the skull-remained and inflammatory-free TIS window provides natural and optimal imaging windows for neuroscience research.

Considering the limitations of previous liquid agents-based skull optical clearing windows in long-term continuous observation and high-resolution monitoring in awake mice, we used three reagents to dissociate collagen, dissolve lipids, perform refractive index matching of the skull, and finally solidify the optical clearing reagent with a coverslip. In this way, the skull could keep transparent. Therefore, the results of *in vitro* experiments and *in vivo* white-light imaging, LSCI and HSI showed that the skull became more and more transparent through each step. The established TIS window could keep the skull transparent for a long time, just like an open-skull glass window. Specifically, the resolving power of cortical blood flow distribution monitoring via LSCI through the TIS window didn’t show a significant decrease over 3 weeks.

Two-photon microscopy is promising in *in vivo* high-resolution sectioning imaging, has been widely used in the applications of research in brain science and related disease models.^27-30^ Our results indicated that the TIS window could remarkably increase the imaging depth and imaging quality of two-photon microscopy, make it possible to track vasculature, axon and dendritic spines in weeks. Long-term monitoring of synaptic dynamics is of great significance in the research of neural science such as motor memories and neuronal circuit establishment, as well as brain dysfunctions such as Parkinson’s disease (PD).^25, 26, 31, 32^ Using two-photon microscopy and thinned-skull window, Xu et al. found that adolescent motor training on mice would enhance spine dynamics within 4 days.^31^ Besides, long-term cortical vascular monitoring is also important in the researches such as aging and stroke.^33, 34^ As a consequence, TIS window assisted long-term two-photon neurovascular tracking provides technical support for basic neuroscience and disease studies. In addition, it was found that our TIS window suits three-photon imaging well, and the neuronal cell bodies at 800 microns, those were unable to be observed by two-photon microscopy, could be distinguished clearly and monitored over 10 days.

Compared to previous optical clearing skull windows, the TIS window showed a big progress. Steinzeig et al. reported a chronic “transparent skull” technique for mice, where they continuously monitored the mouse cortex for 2 months.^35^ However, because of the limited transparency of the treated skull, it was only capable for the very surface observation with limited resolution but didn’t monitor tissue in depths or tiny neural structures. Later, our group developed “SOCW” technique and “USOCA” technique. The SOCW firstly made it possible for mouse synaptic-resolution imaging through intact skull, while it required a little bit thinned skull for adult mice, therefore was limited for repeatable optical clearing and imaging. The USOCA established a switchable optical clearing skull window for repeatable high-resolution cortical observation for a long time. The mice could be optical clearing treated and imaged once a month over 5 months. However, the transparency of the skull gradually decreases with the number of treatments, leading to a limited observation frequency and times. The TIS window developed in this work held advantages of high efficiency of skull optical clearing, no need for skull thinning, and more importantly, during the weeks when the skull remains transparent, the brain could be observed as many times as needed. In the future work, if necessary, one can establish TIS window every few weeks to allow longer continuous observation.

Furthermore, it is the first time that the skull optical clearing window could be used for monitoring fine cortical neural structure and activities in awake mice. In our previous study, we reported skull optical clearing window (SOCW), which was able to perform synaptic two-photon microscopy at the very surface of the cortex, but with the mice anesthetized.^13^ In the SOCW technique, the skull should be cover by glycerinum to keep it transparent. When using a water-immersed objective for two-photon imaging, a plastic wrap needs to be used to separate the glycerin from water. In this case, the interface between glycerol and the plastic wrap, and the interface between plastic wrap and water are easily disturbed by the slight movement of the mice, which has serious negative effects on imaging. On the contrary, using TIS window, the skull, cured S3 and coverslip are relatively still, greatly improving the anti-jamming ability of imaging. Therefore, the clear dendritic images of awake mice were able to be captured.

It is important to evaluate the safety of the TIS window, which aims to determine whether the cortical observation is under a normal state of the brain. In general, the microglia are activated maximally 2 days after craniotomy,^36, 37^ so we carried out *in vivo* and *ex vivo* experiments to observe microglia 2 days after TIS window establishment. In addition, the expression of GFAP is usually upregulated in activated astrocytes 7–14 days after injury,^36, 38^ therefore we visualized GFAP 10 days after TIS window establishment. Furthermore, if there is inflammation in the brain, leukocytes will collect near the area of inflammation. Thus, we performed H&E staining to evaluate if the TIS window would cause leukocyte aggregation. All results indicated that the TIS window technique is safe and won’t lead to any side effects above. These results are consistent with previous reports of non-chronic optical clearing skull windows.^13, 14^

Admittedly, compared to skull removal, the imaging quality in deep tissue through TIS window still need to be improved. After all, optical clearing technique cannot completely eliminate the skull’s attenuation of light. However, this work demonstrated that TIS window was capable enough for certain neurovascular structure and functional observations, indicating its potential in further diagnosis of brain diseases.

In summary, the safe TIS window with easy and quick steps, as a chronic optical clearing skull window, has the advantages of both open-skull glass window and thinned skull window. On one hand, the TIS window allows us to perform cortical imaging immediately after the establishment. On the other hand, the TIS window is compatible with long-term cortical observation. In addition, the TIS window overcomes the disadvantages of the previous optical clearing skull window, which requires repeated operation and is not suitable for the imaging of awake animals. More importantly, the skull-remained TIS window would keep the brain in a normal environment as much as possible. Therefore, the TIS window technique holds great potential for fundamental research in brain science.

## Methods

### Animals

All animal procedures were approved by the Experimental Animal Management Ordinance of Hubei Province, China, and carried out in accordance with the guidelines for the humane care of animals. 8-week-old female *Thy-1-EGFP, Thy-1-YFP* and *Cx3cr1*^*EGFP/+*^ mice were purchased from the Jackson Laboratory (Bar Harbor, ME, USA) and housed and bred in Wuhan National Laboratory for Optoelectronics with a normal cycle (12 h light/dark). 8-week-old female wild-type *BALB/c* and C57 mice were supplied by the Wuhan University Center for Animal Experiment (Wuhan, China). Transgenic mice expressing fluorescent proteins under the control of the Thy1 promoter (Thy1-EGFP and Thy1-YFP) were used for imaging dendritic spines, whereas those expressing the enhanced green fluorescent protein in microglia (Cx3cr1^EGFP/+^) were used for imaging microglia. Wild-type *BALB/c* mice were used for non-stained cerebral blood flow/blood oxygen imaging and FITC-stained vasculature fluorescence imaging. Wild-type C57 mice were used for GCaMP6s-stained calcium imaging.

### Establishment of TIS window

Mice were firstly anesthetized, and a midline incision was made on the scalp along the direction of the sagittal suture. A holder with a hole of 5 mm diameter in the center was glued onto the skull, and the mouse was immobilized in a custom-built plate. The TIS window establishment requested for 3 solutions. First, solution 1 (S1) was applied to the exposed skull for 10 min to dissociate collagen in the skull. After S1 was removed, solution 2 (S2) was then applied to the skull for 5 min to remove lipids in the skull. When S2 was removed and a small quantity of solution 3 (S3) was applied to the skull for RI matching. Then the S3 covered the exposed skull with a thin layer, a 5-mm coverslip was put onto the surface of the S3. Finally, a LED was used to irradiate the skull area for 2 min, making S3 solidify. Thus, the TIS window was established, and cortical observation could be performed either immediately or any time in the future for a long time. The thickness of the solidified S3 was measured to be ∼80 μm.

### In vitro evaluation TIS window technique

Fresh skull samples (0.4×0.4 mm^2^) obtained from the mice were placed in front of a spectrometer (USB4000-VIS-NIR, Ocean Inslight, USA) to evaluate the transmittance enhancement of the skull with the TIS window technique. A broad-spectrum white light source (HL-2000, Mikropack, Germany) was used to measure the transmittance spectrum (400-1000 nm) of the intact skull. Next, the skulls were treated step by step according to the steps used to establish the TIS window, and the change of the transmittance spectrum was recorded (Fig. S2a).

Fresh skulls were taken from mice and cut in half down the middle to macroscopically test the imaging quality through the TIS window. The left parts were subsequently immersed in S1, S2 and S3, followed by a 2-min LED irradiation, while the right parts were used as control. Finally, two parts of the skulls were placed on a grid paper, and a camera was used to take photos (Fig. S2b).

Fresh skull samples (0.4×0.4 mm^2^) obtained from the mice were put on a 1951 United States Air Force (USAF) resolution test target and treated S1, S2, S3 and light irradiation in sequence. A camera (AxioCamHRc, Zeiss, Germany) attached to a stereomicroscope (Zeiss Axio Zoom. V16, Zeiss, Germany) was applied to take the white-light images of skull before and during treatment of the TIS technique to quantificationally decide the increase of imaging resolution. (Fig. S2c).

### Dual-modal imaging system for blood flow/blood oxygen imaging

A home-built dual-modal optical imaging system^39^ was used to obtain blood flow and blood oxygen distribution in the cortical vasculature to *in vivo* evaluate the efficacy of the TIS window. The system consisted of LSCI and HSI. These two imaging modes share a stereoscopic microscope, and each has a light source and a detection system. For LSCI, a He-Ne laser beam went through an adjustable optical attenuator, was used to illuminate the areas of interest after expanded. A CCD camera mounted on a stereomicroscope recorded a sequence of raw speckle images. For HSI, a ring-like LED light with a polarizer was used for illumination. A liquid crystal tunable filter (LCTF, Perkin Elmer, USA) was placed before a CCD camera in another channel to split wavelengths. Laser speckle temporal contrast^40^ and multiple linear regression analysis methods^41^ were used to calculate blood flow and blood oxygen, respectively.

The contrast-to-noise ratio (CNR) was used to access the imaging quality of LSCI and was defined as the following equation:^42^

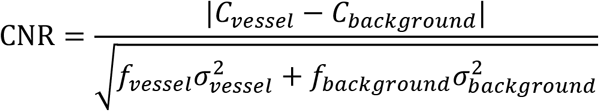

where *C*_*vessel*_ and *C*_*background*_ are the mean contrast values of the blood flow and the background, respectively; 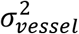 and 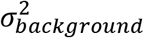 are the variances in the contrast values of the blood flow and the background, respectively; *f*_*vessel*_ and *f*_*background*_ are the fractions of pixels classified as blood flow and the background in all the selected pixels in the speckle contrast image, respectively.

### Two-photon microscopy for cerebrovascular and neural structural imaging

Mice (*Thy-1-EGFP)* with EGFP-expressing dendritic spines/axon/cell body of neurons and FITC-dextran-labeled cerebral vasculature imaging were acquired by a two-photon microscope (Ultima 2p; Bruker, USA) with a Ti:sapphire laser (Chameleon, Coherent, USA) at 900 nm. Mice those were mounted with the holder on the exposed skull were placed under the microscope. Image stacks were captured with a step size of 2 μm utilizing a water-immersed objective (20×, numerical aperture=1.00, working distance = 2 mm, Olympus). Bright-field images of the superficial cortical vasculature were used to relocated and reimaged the region of interest. Although the bright-field imaging quality was limited before skull optical clearing, it is capable to use the low-resolution vasculature map to relocated.

### Two-photon microscopy for calcium imaging in awake mice

Two-photon calcium imaging was performed in the region of somatosensory cortex through the TIS window in awake state. First, a cranial drill was used to drill a small hole into the skull above S1HL (−0.82 mm AP, 1.5 mm ML from bregma)). Second, a glass micropipette and a PicoSprizer III (Parker) were used to inject virus containing GCaMP6s (rAAV-hSyn-Gcamp6s-WPRE-hGH pA, 0.3 μL) into the cortex with an angle of 45° and at depth of 0.3 mm. Next, the TIS window was established after the skull hole was filled up with bone wax. Two weeks later, when the virus was fully expressed, the mouse head with the TIS window was immobilized under the two-photon microscope for imaging, while the paws of the mouse were free to move. Neural calcium imaging was then performed to record electric stimuli activities 100 μm below pia using 2PM with 30 fps. The front-paw electro-stimulation was performed as previously described on the ipsilateral side of the recorded sensorimotor cortex.^43^ The isolated pulse stimulator (AM-system, Model 2100) was used for front-paw electrical stimulation (0.2 mA, 3 Hz, biphasic +/-). The stimulus constant of 8 trials, each trial consists of a 3s baseline period, 3s stimulation events, and 24 s convalescence time for each trial of 30 s.

### Optogenetic experiment in awake mice

Two-photon calcium imaging of optogenetic response was performed in the region of somatosensory cortex through the TIS window in awake state. First, a cranial drill was used to drill a small hole into the skull above S1HL (−0.82 mm AP, 1.5 mm ML from bregma)). Second, a glass micropipette and a PicoSprizer III (Parker) were used to inject virus containing ChrimsonR-tdTomato (rAAV-hSyn-ChrimsonR-tdTomato-WPRE-bGH pA, 0.3 μL) and GCaMP6s (rAAV-hSyn-Gcamp6s-WPRE-hGH pA, 0.3 μL) into the cortex with an angle of 45° and at depth of 0.3 mm. Next, the TIS window was established after the skull hole was filled up with bone wax. Five weeks later, when the virus were fully expressed, the mouse head with the TIS window was immobilized under the two-photon microscope for imaging. Neural calcium imaging was then performed to record optical stimuli activities 100 μm below pia using 2PM with 30 fps. A 590-nm laser (2 mW, emitted from optical fiber) was used as stimulus source and the optical fiber was put above the mouse head. The stimulus constant of 8 trials, each trial consists of a 9 s baseline period, 4 s stimulation events, and 17 s convalescence time for each trial of 30 s.

### Three-photon microscopy imaging

Mice (*Thy-1-YFP)* with YFP-expressing dendritic spines/axon/cell body of neurons were imaged for the suitability of the TIS window in the application of 3PM by long-term monitoring. We used a three-photon microscope (Ultima 2p plus; Bruker, USA) equipped with a water-immersed objective (20×, numerical aperture=1.00, working distance = 2 mm, Olympus), and a 1300 nm fs laser (maximal output power 1.5 W, 400 kHz, 50 fs) from a noncollinear optical parametric amplifier (Spirit-NOPA-VISIR, Spectra Physics, USA) pumped by a regenerative amplifier (Spirit-16, Spectra Physics, USA). The mouse that was mounted with the holder on the exposed skull was placed under the microscope. The image stack was acquired with a depth interval of 5 μm, and pixel dwell time was 2 μs, 4 frames were averaged.

### Safety of TIS window

The *in vivo* and *ex vivo* examinations were performed to explore the activation of microglia cells, the expression of GFAP, and the distribution of neutrophil in the cortex under the TIS window.

The TIS window was established on the left side of the skull of Cx3cr1-GFP mice, in which GFP was expressed in microglia. The mice were then put under the two-photon microscope to monitor microglia for 1 hour. Next, the same region of interest was re-observed *in vivo* 48 hours after window establishment, because microglia activity usually reaches a maximum 48 hours after a craniotomy. After imaging, the mice were perfused and fixed by PBS and 4% paraformaldehyde (PFA, Sigma-Aldrich, USA), then the brain was taken out and placed into 4% PFA overnight for post-fixation. The brain was then sliced (100 μm) and imaged with a confocal microscope (Zeiss, Germany).

Similarly, to test the GFAP expression, we performed the same steps as outlined above, except that the brain slices were immunohistochemically stained with anti-GFAP antibodies 10 days after TIS window establishment, and imaged with the same confocal microscope, since astrocyte activity reaches a maximum at certain time point after a craniotomy.

Since the longest observation in this work last for 21 days, microglia and GFAP expression in were also monitored in brain slices 21 days after TIS window establishment.

Finally, H&E staining was performed onto the brain slices and they were imaged using an upright microscope system (Nikon, Japan), to detect whether neutrophil penetrated the parenchyma, which was a sign of an inflammatory response.

### Data quantification

All imaging data were analyzed with Image J software that was developed by National Institutes of Health (Bethesda, Maryland). 3D image reconstruction was performed in Imaris (Bitplane).

## Author Contributions

Dong-Yu Li was involved in the conceptualization, experiment setup, investigation, statistical analysis, writing and editing. Zheng-Wu Hu was involved in the experiment setup, investigation and statistical analysis. Hequn Zhang was involved in the multi-photon microscope adjustment. Liang Zhu was involved in virus injection. Yin Liu was involved in the optogenetic experiments. Ting-Ting Yu and Jing-Tan Zhu were involved in conceptualization. Wang Xi and Jun Qian was involved in conceptualization, editing and project management. Dan Zhu was involved in conceptualization, writing, editing and project management.

## Competing Interests statement

The authors declare no financial or commercial conflict of interest.

## Acknowledgement

We thank Prof. Tonghui Xu from Fudan University and Dr. Robert Prevedel from European Molecular Biology Laboratory (EMBL) for their advice. This work was supported by National Natural Science Foundation of China (NSFC) (Grant Nos. 61860206009, 81870934, 61975172, 61735016, 91632105, 81961128029, 82001877, 81961138015); the National Key Research and Development Program of China (2017YFA0700501); Fundamental Research Funds for the Central Universities (Nos. 2020-KYY-511108-0007, 2019QNA5001); China Postdoctoral Science Foundation funded project (Nos. BX20190131, 2019M662633); Funding for Postdoctoral Innovation Research Post in Hubei Province and the Innovation Fund of WNLO. The authors also thank to the Optical Bioimaging Core Facility of WNLO-HUST for support in data acquisition.

## Notes

### Competing Interest Statement

The authors have declared no competing interest.

## Reference

1. Laing, B.T., Siemian, J.N., Sarsfield, S. & Aponte, Y. Fluorescence microendoscopy for in vivo deep-brain imaging of neuronal circuits. Journal of Neuroscience Methods 348, 109015 (2021).

2. Calvo-Rodriguez, M., Kharitonova, E.K. & Bacskai, B.J. In vivo brain imaging of mitochondrial Ca2+ in neurodegenerative diseases with multiphoton microscopy. Biochimica et Biophysica Acta (BBA)-Molecular Cell Research, 118998 (2021).

3. Tang, Y. et al. In vivo two-photon calcium imaging in dendrites of rabies virus-labeled v1 corticothalamic neurons. Neuroscience bulletin 36, 545–553 (2020).

4. Ji, N., Freeman, J. & Smith, S.L. Technologies for imaging neural activity in large volumes. Nat Neurosci 19, 1154–1164 (2016).

5. Hong, G.S., Antaris, A.L. & Dai, H.J. Near-infrared fluorophores for biomedical imaging. Nat Biomed Eng 1, 0010 (2017).

6. Fan, J.L. et al. High-speed volumetric two-photon fluorescence imaging of neurovascular dynamics. Nat Commun 11, 6020 (2020).

7. Kisler, K. et al. In vivo imaging and analysis of cerebrovascular hemodynamic responses and tissue oxygenation in the mouse brain. Nat Protoc 13, 1377–1402 (2018).

8. Kneipp, M. et al. Effects of the murine skull in optoacoustic brain microscopy. J Biophotonics 9, 117–123 (2016).

9. Fan, X.F., Zheng, W.T. & Singh, D.J. Light scattering and surface plasmons on small spherical particles. Light-Sci Appl 3, e179 (2014).

10. Holtmaat, A. et al. Long-term, high-resolution imaging in the mouse neocortex through a chronic cranial window. Nat Protoc 4, 1128–1144 (2009).

11. Yang, G., Pan, F., Parkhurst, C.N., Grutzendler, J. & Gan, W.B. Thinned-skull cranial window technique for long-term imaging of the cortex in live mice. Nat Protoc 5, 201–208 (2010).

12. Dorand, R.D., Barkauskas, D.S., Evans, T.A., Petrosiute, A. & Huang, A.Y. Comparison of intravital thinned skull and cranial window approaches to study CNS immunobiology in the mouse cortex. Intravital 3, e21978 (2014).

13. Zhao, Y.J. et al. Skull optical clearing window for in vivo imaging of the mouse cortex at synaptic resolution. Light-Sci Appl 7, 17153 (2018).

14. Zhang, C. et al. A large, switchable optical clearing skull window for cerebrovascular imaging. Theranostics 8, 2696–2708 (2018).

15. Zhang, C. et al. Photodynamic opening of the blood-brain barrier to high weight molecules and liposomes through an optical clearing skull window. Biomed Opt Express 9, 4850–4862 (2018).

16. Zhang, C. et al. Age differences in photodynamic therapy-mediated opening of the blood-brain barrier through the optical clearing skull window in mice. Laser Surg Med 51, 625–633 (2019).

17. Feng, W. et al. Comparison of cerebral and cutaneous microvascular dysfunction with the development of type 1 diabetes Theranostics 9, 5854–5868 (2019).

18. Wang, J., Zhang, Y., Xu, T.H., Luo, Q.M. & Zhu, D. An innovative transparent cranial window based on skull optical clearing. Laser Phys Lett 9, 469–473 (2012).

19. Hirshburg, J.M., Ravikumar, K.M., Hwang, W. & Yeh, A.T. Molecular basis for optical clearing of collagenous tissues. J Biomed Opt 15, 055002 (2010).

20. Chen, Y.G. et al. Coherent Raman Scattering Unravelling Mechanisms Underlying Skull Optical Clearing for Through-Skull Brain Imaging. Anal Chem 91, 9371–9375 (2019).

21. Yang, B. et al. Single-Cell Phenotyping within Transparent Intact Tissue through Whole-Body Clearing. Cell 158, 945–958 (2014).

22. Wang, T.Y. et al. Three-photon imaging of mouse brain structure and function through the intact skull. Nat Methods 15, 789-+ (2018).

23. Horton, N.G. et al. In vivo three-photon microscopy of subcortical structures within an intact mouse brain. Nat Photonics 7, 205–209 (2013).

24. Li, D.Y. et al. Visible-near infrared-II skull optical clearing window for in vivo cortical vasculature imaging and targeted manipulation. J Biophotonics 13, e202000142 (2020).

25. Guo, L. et al. Dynamic rewiring of neural circuits in the motor cortex in mouse models of Parkinson’s disease. Nat Neurosci 18, 1299–1309 (2015).

26. Xu, T., Wang, S., Lalchandani, R.R. & Ding, J.B. Motor learning in animal models of Parkinson’s disease: Aberrant synaptic plasticity in the motor cortex. Mov Disord 32, 487–497 (2017).

27. Qin, Z. et al. Adaptive optics two-photon endomicroscopy enables deep-brain imaging at synaptic resolution over large volumes. Science advances 6, eabc6521 (2020).

28. Fan, J.L. et al. High-speed volumetric two-photon fluorescence imaging of neurovascular dynamics. Nature communications 11, 1–12 (2020).

29. Qi, J. et al. Aggregation-induced emission luminogen with near-infrared-II excitation and near-infrared-I emission for ultradeep intravital two-photon microscopy. Acs Nano 12, 7936–7945 (2018).

30. Chen, C.P. et al. In Vivo Near-Infrared Two-Photon Imaging of Amyloid Plaques in Deep Brain of Alzheimer’s Disease Mouse Model. Acs Chem Neurosci 9, 3128–3136 (2018).

31. Xu, T. et al. Rapid formation and selective stabilization of synapses for enduring motor memories. Nature 462, 915–919 (2009).

32. Yu, X. et al. Accelerated experience-dependent pruning of cortical synapses in ephrin-A2 knockout mice. Neuron 80, 64–71 (2013).

33. Zaorsky, N.G., Zhang, Y., Tchelebi, L.T., Mackley, H.B. & Zacharia, B.E. Stroke among cancer patients. Nature Communications 10 (2019).

34. Peters Ageing and the brain. Postgraduate Medical Journal 82, 84–88 (2006).

35. Steinzeig, A., Molotkov, D. & Castren, E. Chronic imaging through “transparent skull” in mice. Plos One 12 (2017).

36. Drew, P.J. et al. Chronic optical access through a polished and reinforced thinned skull. Nat Methods 7, 981–U960 (2010).

37. Xu, H.T., Pan, F., Yang, G. & Gan, W.B. Choice of cranial window type for in vivo imaging affects dendritic spine turnover in the cortex. Nat Neurosci 10, 549–551 (2007).

38. Zuo, Y., Lin, A., Chang, P. & Gan, W.B. Development of long-term dendritic spine stability in diverse regions of cerebral cortex. Neuron 46, 181–189 (2005).

39. Feng, W. et al. Comparison of cerebral and cutaneous microvascular dysfunction with the development of type 1 diabetes. Theranostics 9, 5854–5868 (2019).

40. Draijer, M., Hondebrink, E., Leeuwen, T.V. & Steenbergen, W. Review of laser speckle contrast techniques for visualizing tissue perfusion. Lasers in Medical Science 24, 639 (2009).

41. Wei, F., Rui, S., Chao, Z., Yu, T. & Picker, O. Lookup-table-based inverse model for mapping oxygen concentration of cutaneous microvessels using hyperspectral imaging. Optics Express 25, 3481–3495 (2017).

42. Wang, J., Zhang, Y., Li, P., Luo, Q. & Zhu, D. Tissue optical clearing window for blood flow monitoring. IEEE Journal of Selected Topics in Quantum Electronics 20, 92–103 (2013).

43. Winship, I.R. & Murphy, T.H. In vivo calcium imaging reveals functional rewiring of single somatosensory neurons after stroke. J Neurosci 28, 6592–6606 (2008).

